# The genomic basis of evolved virus resistance is dependent on environmental resources

**DOI:** 10.1101/666404

**Authors:** Katherine Roberts, Sean Meaden, Stephen Sharpe, Suzanne Kay, Toby Doyle, Drew Wilson, Lewis J. Bartlett, Steve Paterson, Mike Boots

## Abstract

Parasites impose strong selection on their hosts, but the level of resistance evolved may be constrained by the availability of resources. However, studies identifying the genomic basis of such resource mediated selection are rare, particularly in non-model organisms. Here, we investigated the role of nutrition in the evolution of resistance to a DNA virus (PiGV), and associated trade-offs, in a lepidopteran pest species (*Plodia interpunctella*). Through selection experiments and whole genome sequencing we identify putative mechanisms of resistance that depend on the nutritional environment during selection. We find that the evolution of resistance is specific to diet, with adaptation to a low nutrition diet constraining resistance when challenged with the pathogen on a high nutrition diet. Resistance in a low nutrition environment is negatively correlated with growth rate, consistent with an established trade-off between immunity and development. Whole genome resequencing of the host shows that resistance mechanisms are highly polygenic and suggests evidence for trade-offs at the genetic level. Critically when populations evolve in high resource conditions, resistance is linked to metabolic and immune pathways, however it is more closely associated with cytoskeleton organisation when selected under low nutrition. Our results emphasise the importance of resources on the evolution of resistance.

## Introduction

Parasites and pathogens impose strong selection on their hosts resulting in the evolution of a range of defence mechanisms. For example, invertebrates possess an effective innate immune system that is capable of fighting infections from a wide range of pathogens (Kingsolver *et al*. 2013, Sackton *et al.* 2007, Viljakainen *et al.* 2015). Hosts can also use a range of pathways to prevent or tolerate infection, including behavioural or physiological changes (Raberg *et al.* 2009; Curtis *et al.* 2011, Lefevre et al. 2012). The host strategy most likely to evolve, or be maintained, will ultimately depend upon the resources available to the host, as resistance mechanisms are both costly to initiate and maintain (Cotter, Simpson, Raubenheimer, & Wilson, 2011; Knutie, Wilkinson, Wu, Ortega, & Rohr, 2017; Lochmiller & Deerenberg, 2000). Such resource availability can vary due to both temporal and spatial differences e.g. seasonality, population density and patchiness of resource availability, and in terms of the quantity and quality of required resources. A core component of resource availability is nutrition, which is likely to greatly impact the evolution of resistance to parasites. For example, greater resistance is predicted to evolve under higher resource environments for two reasons. Firstly, reduced competition for resources should allow organisms to invest more in resistance mechanisms. Secondly, higher resources can lead to greater population density and therefore greater transmission events and chance of infection, resulting in stronger selection for resistance (Lopez-Pascua & Buckling 2008; Gómez *et al.* 2015). Resistance mechanisms may also, in principle, be specific to host nutritional status, where a resource threshold is required for a resistance mechanism to be induced and functionally useful.

Such resistance mechanisms typically come at a price: either through the activation of induced defence mechanisms (Graham, Allen, & Read, 2005; Moret & Schmid-Hempel, 2000; Sadd & Siva-Jothy, 2006) or through the maintenance of a constitutively expressed defence when parasites are absent (Boots & Begon, 1993; Fuxa & Richter, 1992; Kraaijeveld & Godfray, 1997; McKean, Yourth, Lazzaro, & Clark, 2008). Such costs may lead to the stable maintenance of polymorphism within a population (Antonovics & Thrall 1994; Juneja & Lazzaro 2009, Bowers, Boots & Begon 1994, Boots and Haraguchi 1999). Therefore, understanding how such mechanisms evolve, or are maintained, in controlled laboratory conditions is important for predicting the evolution of resistance in more variable wild populations. Here we examined the role of nutrition in the evolution of resistance to a DNA virus in its insect host in response to natural oral infection using an experimental evolution approach. We use the Indian Meal Moth, *Plodia interpunctella* and its naturally occurring granulosis baculovirus (PiGV) as a model system, where we have previously demonstrated that there is a resource-dependent cost to the evolution of resistance (Boots, 2011). Both the level of resistance attained and the associated costs depended on the selection environment, suggesting that different resistance mechanisms may be forced to evolve in different environments (Boots, 2011). We therefore evolve populations for multiple generations on two different nutritional environments, either in the presence or absence of the viral pathogen. We then tested the strength of the populations’ resistances to the viral pathogen, and the larval development across the nutritional levels in order to quantify any potential trade-offs. Finally, we used whole-genome resequencing of populations to perform a genome scan for candidate resistance genes. Resequencing experiments are a powerful tool to identify the genetic basis of observed traits, including those involved in host-parasite interactions (Eoche-Bosy et al., 2017; Martins et al., 2014). To date many studies rely on model systems such as *Drosophila* and critically use only a single, often very high nutritional quality diet to identify the genetic basis of variance in traits of interest (Jha et al., 2015; Michalak, Kang, Sarup, Schou, & Loeschcke, 2017; Shahrestani et al., 2017; Turner & Miller, 2012). By studying multiple nutritional levels in a non-model organism we aim to tease apart the contribution of diet to resistance and any relevant trade-offs in a broader context.

## Materials and Methods

### Selection Experiment

Replicate selection lines of the Indian Meal Moth, *Plodia interpunctella* were set up at two different nutritional quantities, both in the presence and absence of a natural pathogen, the granulosis baculovirus PiGV. In order to establish genetically diverse and homogenous selection lines, we initially established a large outbred population of *Plodia interpunctella* by outcrossing existing laboratory strains with a number of populations received from the USDA. The initial set up of selection was based on the methods of (Boots, 2011). Briefly, for the virus selection lines PiGV was mixed into the food medium in which moths both feed on and reproduce within. Larvae become orally infected, which is the natural route of infection through ingesting the infective viral particles whilst they feed. There is therefore a strong selection pressure on all larvae across instar stages.

The resource-level quality of the moth’s food is precisely controlled by the addition of methyl cellulose (an indigestible bulking agent) to the medium (Boots & Begon, 1994). The resource levels to establish our selection lines were determined based on (Boots, 2011). The basic food consisted of a cereal base (50% Ready Brek ©, 30% wheat bran, and 20% ground rice), brewer’s yeast, honey, and glycerol (see supplementary for full methods for resources). To produce the two selection line food levels 10% of the mix was replaced with methyl cellulose (MC) to give the high-quality resource level, and 55% food mix was replaced with MC for the low-quality diet.

Initially, 4 control (no virus) and 4 virus populations were established, on each of the two food resources. Sixty, 3-day post eclosion moths of mixed sex were placed in a 500ml Nalgene pot using an excess of each food mixture (200g). These 16 populations (4 × Control-Low food, 4 × Virus-Low food, 4 × Control-High food and 4 × Virus-High food) constituted one block of the experiment. This set up was repeated for 5 replicate blocks to give 20 separate populations of each of the potential selection regimes. All populations were maintained in incubators at 27°C, 16 Light:8 Dark cycle, and pots were rotated around the incubator in order to control of any effects of incubator position. The day of first eclosion of each pot was noted, and three days post this first eclosion moths were moved onto the next generation as a way controlling for the effect of food on developmental time and ensuring the median day of eclosion was always used to generate the parents of the next generation (Boots, 2011). The populations were maintained for 12 generations in this manner after which they were assayed for their viral resistance and life history traits.

### Phenotypic assays

After 12 generations all populations were relaxed from their selection regime. Populations were split onto two different food types; a high quality 0% MC food (common garden environment i.e. both high and low nutrition treatment populations were reared and assayed on a common diet with no addition of MC) or the food type that they had been selected on for the course of the experiment (10% MC or 55% MC). None of the food for this “relaxed” generation contained any virus but the population set up was otherwise the same as for the selection regime. From these populations, second-generation (to avoid maternal effects), third-instar larvae were either bioassayed with a viral solution, to look at virus resistance, or allowed to develop individually for life history measures. Both assays were carried out in on individuals housed in a segmented 25 well petri dish with an excess of relaxed generation food. The infection assay followed the protocol of Boots & Roberts (2012) where third instar larvae were removed from each population and starved for two hours before being orally dosed with a freshly prepared virus solution diluted with distilled water, 0.1% Coomassie Brilliant Blue R dye (ingestion is indicated by the presence of blue dye in the gut) and 2% sucrose (to encourage feeding). For this experiment, each of the relaxed populations was dosed at 5 different virus concentrations, highest dose of 2.5×10^-4^ % virus solution to dye solution, with four further 1:10 dilutions. A control solution of the blue dye, sucrose solution was also used as a control for dosing protocol. Approximately 25 larvae were dosed at each dose of virus, from each population. Larvae were kept in incubators and the numbers of subsequently infected larvae were recorded as a binary response on visual inspection infected larvae are clearly visible because of their opaque white colour due to a build-up of viral occlusion bodies. As PiGV is an obligate killer, there is no tolerance to infection. We therefore refer to resistance here as the proportion of individuals surviving following viral challenge.

At the same time as the larvae for the infection assay were collected 25 larvae were again individually placed into the 25 well Petri dishes containing high quality resource and allowed to develop in standard incubators conditions. The time to pupation was checked daily and the day that a brown pupa was seen it was removed from its silk case and weighted and recorded.

### DNA extraction and Sequencing

Our methods for studying the genetic basis of PiGV resistance was to use a ‘pool-seq’ approach where individual larvae from a population are pooled and the subsequent extracted DNA is sequenced to generate estimates of allele frequencies within a population. This approach has been developed and validated in a number of papers (Kofler, Langmüller, Nouhaud, Otte, & Schlötterer, 2016; C Schlötterer et al., 2015; Christian Schlötterer, Tobler, Kofler, & Nolte, 2014) and is an efficient way of comparing large numbers of populations. Genomic DNA from each population was extracted using a Blood and Tissue DNA extraction kit (Qiagen, UK). First, 50 larvae from each population were fully homogenised in ATL lysis buffer, and after Proteinase-K digestion the max volume for the column was taken through for the rest of the extraction protocol. (25mg tissue was the max for the column and 180uL of lysate equated to 25mg of original tissue). In parallel, DNA was extracted from 8 individual larvae in order to generate a high confidence SNP dataset, using the QIAGEN Genomic-tip 20/G standard protocol (Qiagen, UK). All samples were sequenced at the University of Liverpool from Illumina TruSeq Nano libraries with 350bp inserts using 125bp paired-end reads on an Illumina HiSeq2500 platform. Reads were quality filtered to remove adapter sequences, reads shorter than 10bp and reads with a minimum window score of 20 using cutadapt (version 1.2.1) (Martin, 2011) and Sickle (version 1.2 (Joshi & Fass, 2011)). Reads were mapped to the *Plodia interpunctella* reference genome (described here and available from LepBase.org) using Bowtie2. GATK’s HaplotypeCaller program was used to generate high confidence SNP markers from sequences obtained from individual larvae. Allele frequency counts were filtered to exclude SNPs with coverage greater than the median plus 3 standard deviations in order to exclude sequencing errors that could occur from mapping to collapsed repeats in the assembly. This SNP dataset was used as a reference dataset to generate allele frequencies at each marker per population using the pool-seq data using Samtools mpileup. Sequence data has been deposited in the ENA under accession number PRJEB27964. Additional sequencing was undertaken to improve the scaffold lengths of the assembly using a proximity ligation method at Dovetail Genomics (Santa Cruz, CA, USA). This method creates chromatin cross links on input DNA, followed by proximity ligation to mark the physical proximity of sequences to each other (Putnam et al., 2016).

### *De novo* assembly and annotation of the *P. interpunctella* genome

In order to reduce heterozygosity prior to genome assembly, a line of *Plodia* was generated by full-sib matings for 10 generations. DNA was extracted using Qiagen GenomeTip and used to make Illumina TruSeq PCR-free, paired-end libraries with insert sizes of c. 350bp, 450bp and 600bp and sequenced on an Illumina MiSeq platform to generate c. 18Gbp of 2×250bp reads and c. 40Gbp of 2×100bp reads on the Illumina HiSeq2000 platform. Nextera mate-pair libraries with 3Kbp and 10Kbp insert sizes were sequenced on the Illumina 2000 platform with c. 50m pairs of 100bp reads from each library. Illumina polyA-ScriptSeq RNA libraries were prepared from 15 individuals and sequenced on the Illumina 2000 platform with c. 45m pairs of 100bp reads from each library. Illumina MiSeq reads were trimmed to Q≥30 and adaptors removed using Sickle and Perl and assembled using Newbler (Roche GS-Assembler v2.6) with flags set for large genome and a heterozygote sample. Mate-pair reads were first mapped to these contigs using Bowtie2 (Langmead & Salzberg, 2012) to remove duplicates and wrongly orientated reads, and scaffolded into contigs using SSPACE (Boetzer, Henkel, Jansen, Butler, & Pirovano, 2011). Gap filling was achieved using GapFiller for 2× 250bp and 2× 100bp paired-end reads and run for three iterations. RNAseq data were mapped to scaffolds within the assembled genome greater than 3Kbp using TopHat2 to identify transcribed regions and splice junctions. These, together with RNAseq data assembled using Trinity, were passed to the MAKER pipeline (Cantarel et al., 2008) to predict genes.

### Phenotypic data analysis

To test the role of diet and exposure to PiGV resistance we used a linear mixed effect model. We used the proportion of surviving larvae at the median assay dose as the response term as this dosage exhibited the largest variance (0.042, Fig. 1A). All selection lines were assayed in both the common garden diet (with no MC replacement) or their respective selection diet (10% or 55% replacement). Selection treatment (PiGV exposure vs. control), assay diet (common garden vs. home) and selection diet (high vs. low) and interactions among these variables were fitted as fixed effects with block (population start date) included as a random effect and a binomial error structure applied. Checks of model residuals showed that the data conform to model assumptions. ANOVA was used to determine p-values following model simplification using AIC. Post-hoc comparisons were made using Tukey’s all-pair comparison with p-values adjusted using the Holm-Bonferroni method. For the developmental data, a similarly structured model was applied, in this case including resistance as a fixed effect and mean growth rate (per population) as the response term.

**Figure 1.**
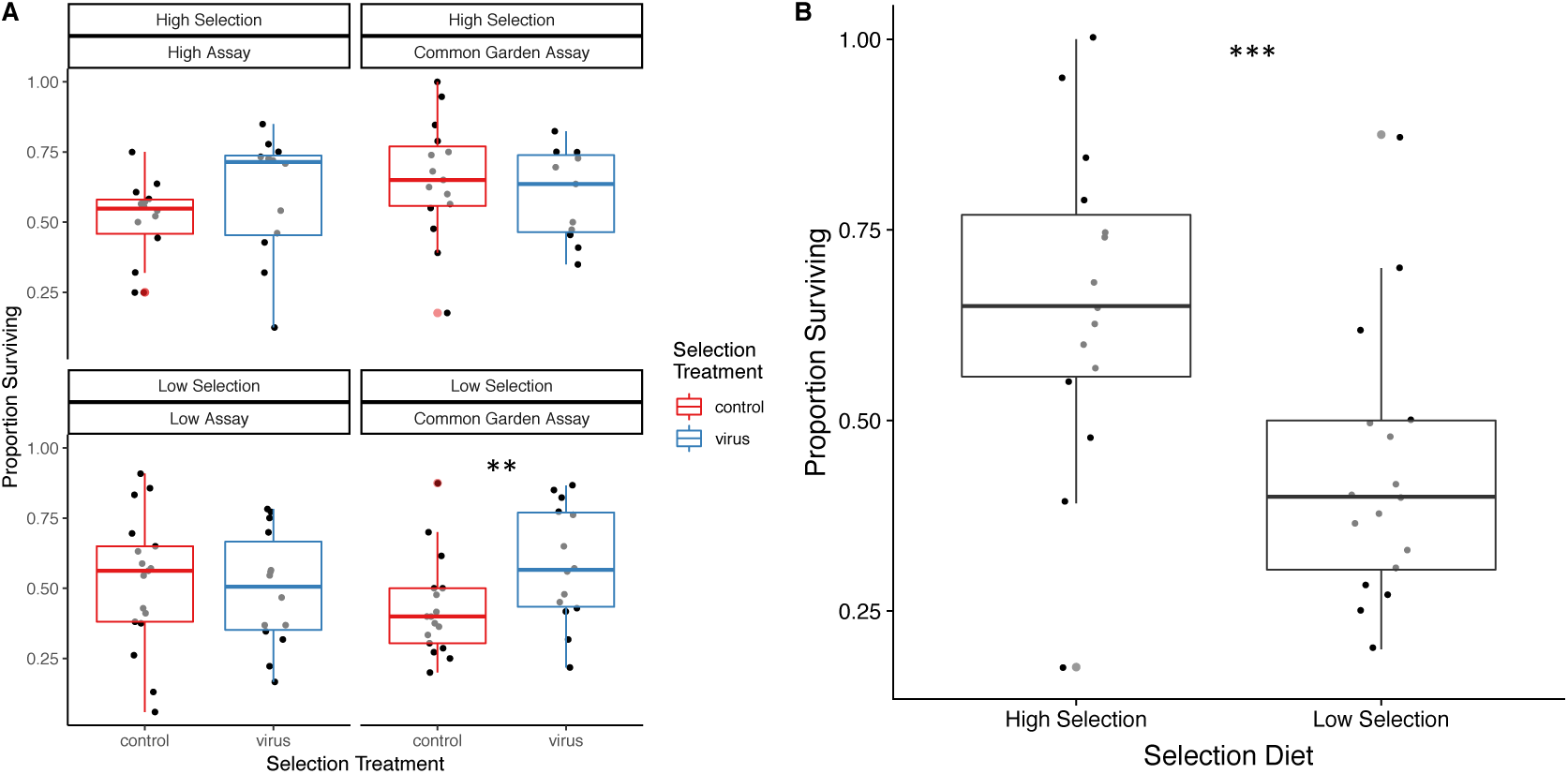
A) Resistance of larval populations when assayed on the diet they were selected on, high nutrition or low nutrition (selected), or a common garden diet. B) Resistance of control populations on their selected diets at the median dose of PiGV.

### Pool-seq genome wide association test

Association tests were run by iterating a binomial GLM on each SNP marker using the alternative and reference allele count as the response variable and the proportion of surviving larvae as the explanatory variable. P-values were computed using stepwise ANOVA. Any SNPs where a model failed to converge or that resulted in a regression containing data points with a Cook’s distance greater than 1 were discarded. This resulted in a filtered dataset of ∼450,000 and 250,000 SNPs (common garden diet and low nutrition respectively). P-values were corrected for a false discovery rate using the Benjamini and Hochberg correction and plotted across the length of the genome to identify regions associated with PiGV resistance. Genetic structure was assessed with the program Baypass using subsets of the data (49686 markers per group, 20 groups total) to assess FMD statistics (distance between covariance matrices, (Förstner & Moonen, 2003).

### Putative function analysis

Functional analysis of the candidate loci resulting from the association tests was conducted in two ways. Firstly, all genes containing associated SNPs were extracted and linked to the *Plodia interpunctella* predicted gene set. In this case significance was defined as P < 0.0001 after false discovery rate correction (Benjamini-Hochberg method) in order to reduce spurious matches. Orthologous genes between *P. interpunctella* and *Drosophila melanogaster* were identified using InParanoid (version 4.1). The resulting UniProt codes from matched genes were used for gene set enrichment analysis using the AmiGO service (http://amigo.geneontology.org/amigo) using Fisher’s exact test. A second approach was to search the BLAST database directly for the best hits to the *P. interpunctella* genes of interested. The resulting best hits were extracted and used to search the UniProt database for gene ontology terms and functional characterizations.

## Results

### Evolution of resistance to PiGV is diet dependent

By comparing virus (PiGV) exposed and unexposed controls selection treatments across multiple diets, we can test a number of specific hypotheses regarding the role of nutrition on resistance evolution. For example, comparing exposed and unexposed controls on their local diet (high or low nutrition) tests whether resistance evolved during the selection experiment. Testing these same populations on a common garden diet tests whether any evolved resistance mechanisms work across environments. A comparison of unexposed controls on a common garden diet allows us to test an effect of diet itself in shaping resistance. Finally, comparing exposed populations selected on a high or low nutrition diet and assayed on a common garden diet tests whether there is a difference in effectiveness between diet-specific resistance mechanisms.

In order to answer these questions, we first assessed resistance to PiGV across all diets and a gradient of doses in order to identify the dosage that maximized variation (Fig. 1A, stock dilution of 2.5e-06). At this dosage we found a strong three-way interaction between the selection diet, assay diet and the selection treatment (exposure to the virus vs. control) on resistance (GLMM, 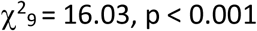, Fig. 1B). To understand the drivers of this effect we used post-hoc testing to compare survivorship among the contrasts that test the hypotheses outlined above (Table 1).

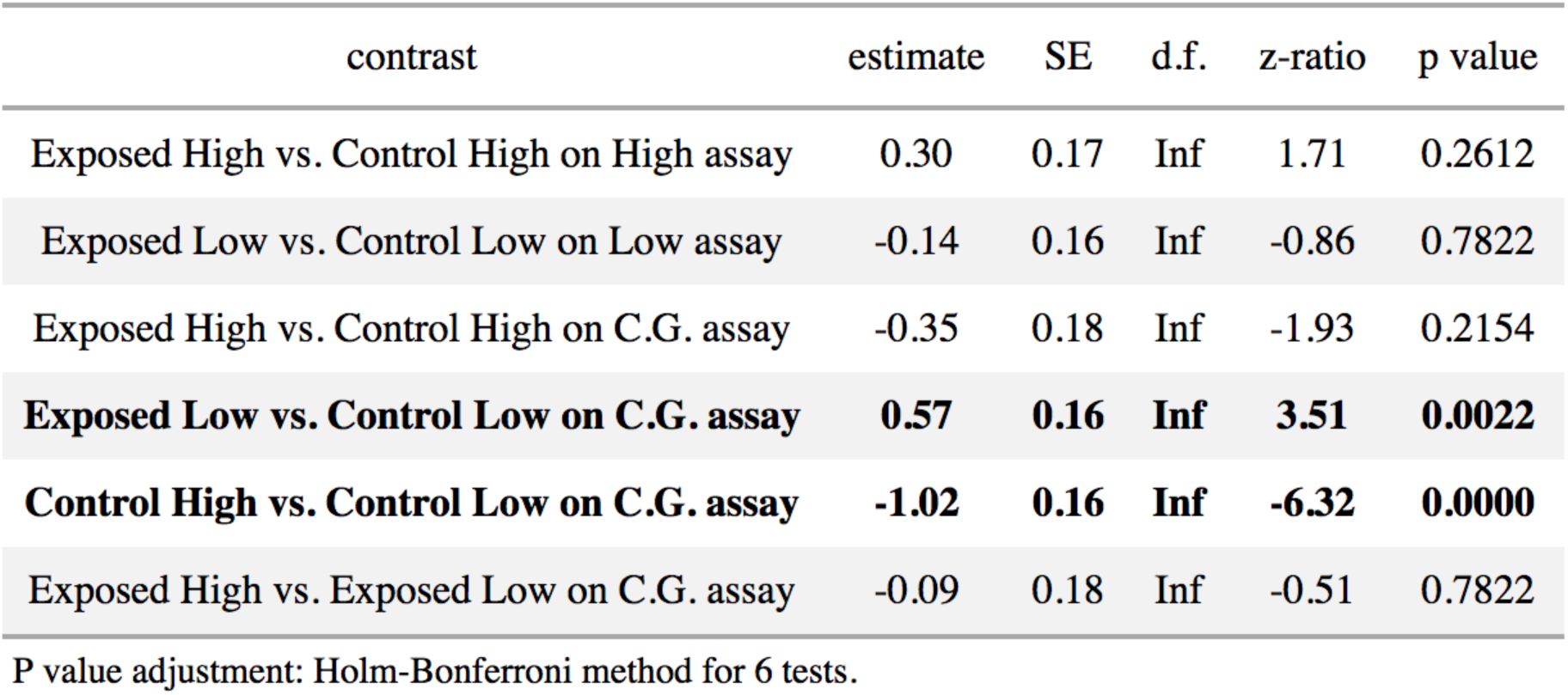

| contrast | estimate | SE | d.f. | z-ratio | p value |
| --- | --- | --- | --- | --- | --- |
| Exposed High vs. Control High on High assay | 0.30 | 0.17 | Inf | 1.71 | 0.2612 |
| Exposed Low vs. Control Low on Low assay | -0.14 | 0.16 | Inf | -0.86 | 0.7822 |
| Exposed High vs. Control High on C.G. assay | -0.35 | 0.18 | Inf | -1.93 | 0.2154 |
| **Exposed Low vs. Control Low on C.G. assay** | **0.57** | **0.16** | **Inf** | **3.51** | **0.0022** |
| **Control High vs. Control Low on C.G. assay** | **-1.02** | **0.16** | **Inf** | **-6.32** | **0.0000** |
| Exposed High vs. Exposed Low on C.G. assay | -0.09 | 0.18 | Inf | -0.51 | 0.7822 |

P value adjustment: Holm-Bonferroni method for 6 tests.

We found the largest effect to be driven by diet itself, for example when comparing just unexposed control populations we find that those selected on a high nutrition diet produce larvae more likely to survive when facing viral challenge on a common garden diet (Table 1.; Fig. 1). This is interesting as it suggests a trade-off between being able to survive in a nutritionally limited environment and resistance to a parasite.

Secondly, we found that larvae from populations exposed to PiGV and selected on a low nutrition diet showed greater survival than their counterpart controls, but only when assayed on the common garden diet (Fig 1, panel A). This suggests that the method of resistance being employed by populations evolved in the two environments is different across the environments. It is interesting to note that we did not observe any difference between larvae from exposed and unexposed populations who were selected on a low nutrition environment, when we assayed them on nutrient limited food. Yet the clear difference on the common garden diet suggests there is selection on resistance between these populations. We also did not identify any differences between exposed and control populations from the high nutrition treatment (Fig. 1, panel A). This suggests that the differences seen in the low nutrition treatments may potentially be the result of a loss of a costly resistance mechanism that is non-functional in a low nutrient environment.

### Diet determines developmental trade-offs

To assess the effects of diet and PiGV exposure on growth rate we assayed all populations on the diet they were evolved on and a common garden diet, as in the resistance assays. We found no significant interactions between assay diet, selection diet, and population exposure, but did find significant independent effects of assay diet and population exposure (i.e. no effect of selection diet) (Fig. 2). Assay diet had the largest effect on growth rate with the fastest growth rates occurring on the common garden diet compared to selected diets (χ^2^ = 157.06, df = 6, p < 0.0001). There was also a significant effect of PiGV exposure during selection, with populations selected for virus resistance exhibiting quicker growth rates than control populations (χ^2^ = 5.05, df = 6, p < 0.025), which is counter to the trade-off observed previously in this system (Boots, 2011).

**Figure 2.**
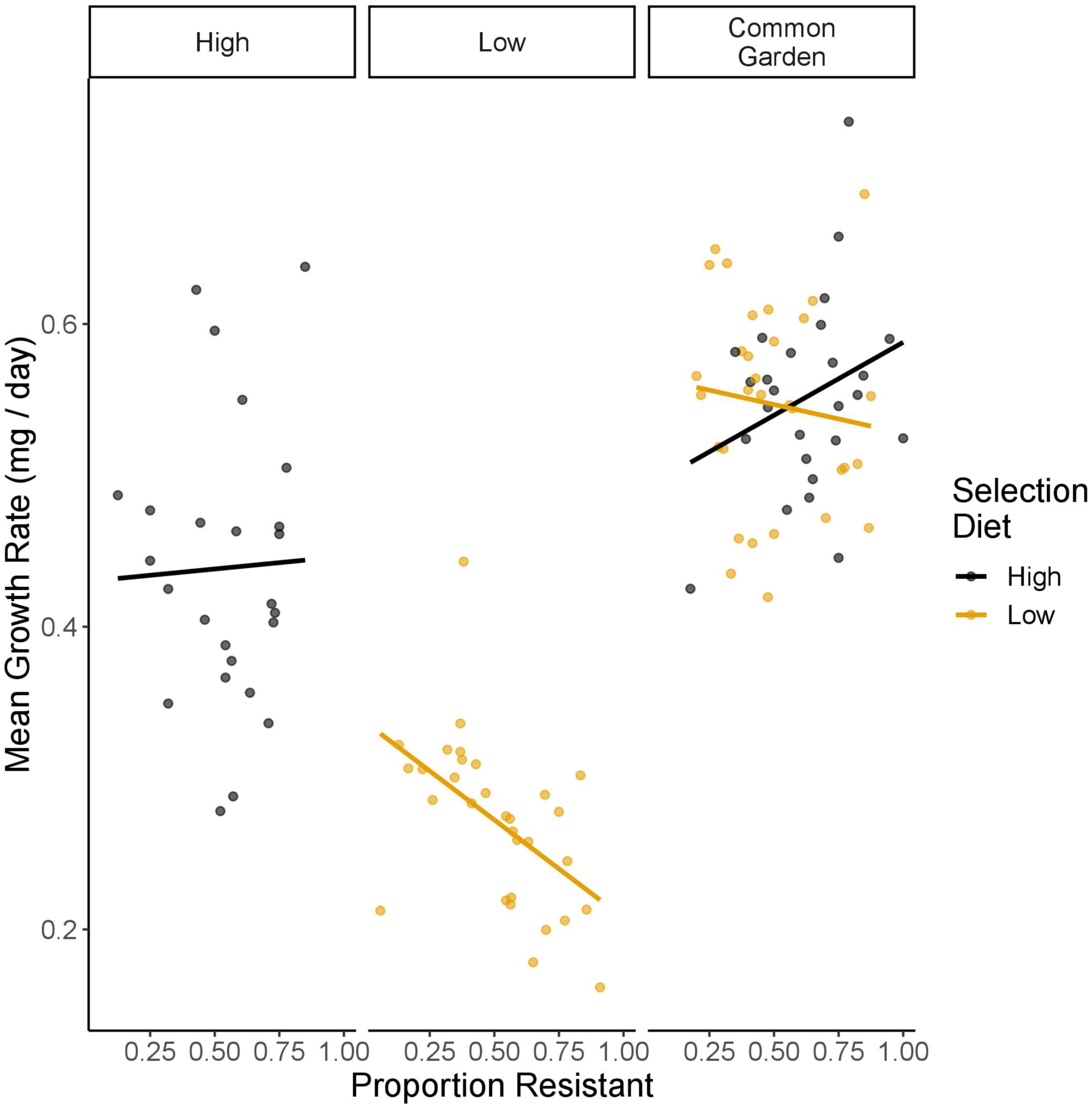
Nutrient dependent trade-offs between mean population growth rate and resistance under all diets. Points represent raw data and lines are model predictions. Yellow points denote larval populations selected on a low-nutrition diet, whilst black denote populations selected on a high nutrition diet. Both resistance and growth rate are population level estimates based on either survivorship or development assays of 25 larvae per population.

As we observed wide variation in growth rates within selection treatments we correlated the mean growth rate of each population to resistance on either their selected or common garden diet irrespective of selection treatment. In this case, we found a strong negative correlation between resistance and growth rate for the low nutrition populations on their selected diet (χ^2^ = 7.35, df = 6, p = 0.0067, Fig.2), but no directional correlation for the high nutrition populations on their selected diet (χ^2^ = 0.028, df = 6, p = 0.866, Fig. 2). On the common garden diet we found a significant interaction between measured resistance and selection diet (high or low nutrition) as predictors of growth rate (χ^2^ = 4.72, df = 7, p = 0.0298, Fig. 2). This demonstrates that the nutritional environment larvae are selected on leads to fundamentally different costs of resistance. In this experiment low nutrient environments selected for a form of resistance that is traded off with growth rate, whereas in the high nutrient environment selection for resistance came at no cost to growth rate.

### Identifying the genomic basis of resistance

To identify the genomic basis of resistance to PiGV infection we employed a genome wide scan in order to associate specific loci with resistance. This method tests the association between allele frequency at each SNP present in our dataset and the resistance of each population. We first assessed SNPs that predicted resistance on a common dietary background i.e. the proportion of surviving larvae when assayed on the common garden diet. We identified a number of candidate SNPs that were strongly correlated with resistance (Fig. 3). Whilst these SNP markers were strongly correlated with PiGV resistance-required by the association test used, they were not correlated with development.

**Fig 3.**
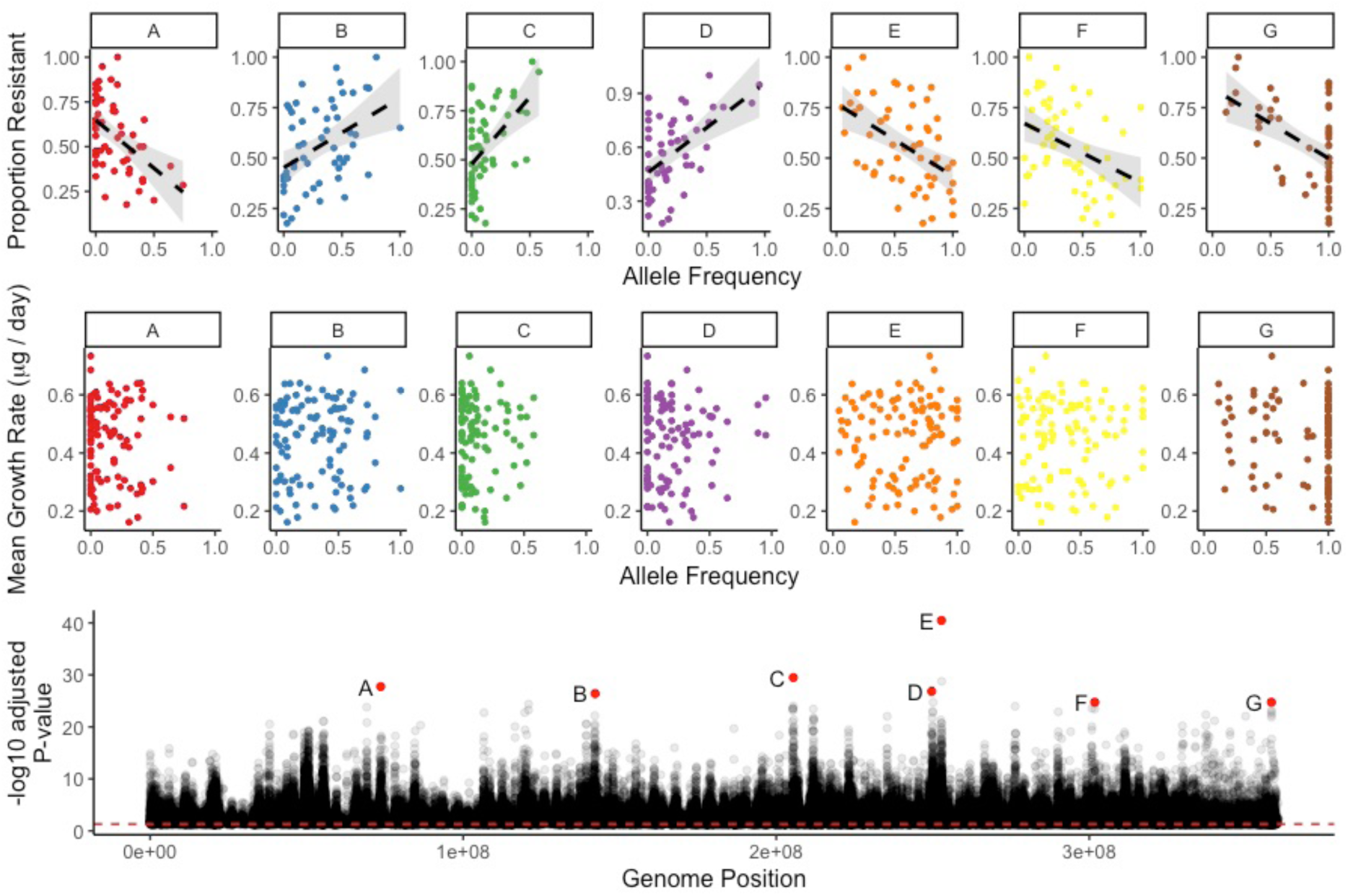
Whole genome scan for SNPs associated with PiGV resistance, regardless of diet (bottom panel). By definition of the methods used for the association test, the allele frequency of these SNPs must be correlated with resistance (top panel). No correlations are seen between allele frequency and growth rate (middle panel).

Whilst assessing the correlations of individual SNPs is useful for identifying strong effect loci, identifying such a large number of SNPs across many scaffolds is suggestive of a polygenic trait and therefore enrichment analyses may be more appropriate for functional inference. Following gene set enrichment analysis, we found that the gene ontology term most overrepresented was calcium ion homeostasis (15-fold enrichment). Neuromuscular synaptic transmission and immune response pathways were also overrepresented and showed 8 and 7-fold enrichments respectively (Fig. 4).

**Figure 4.**
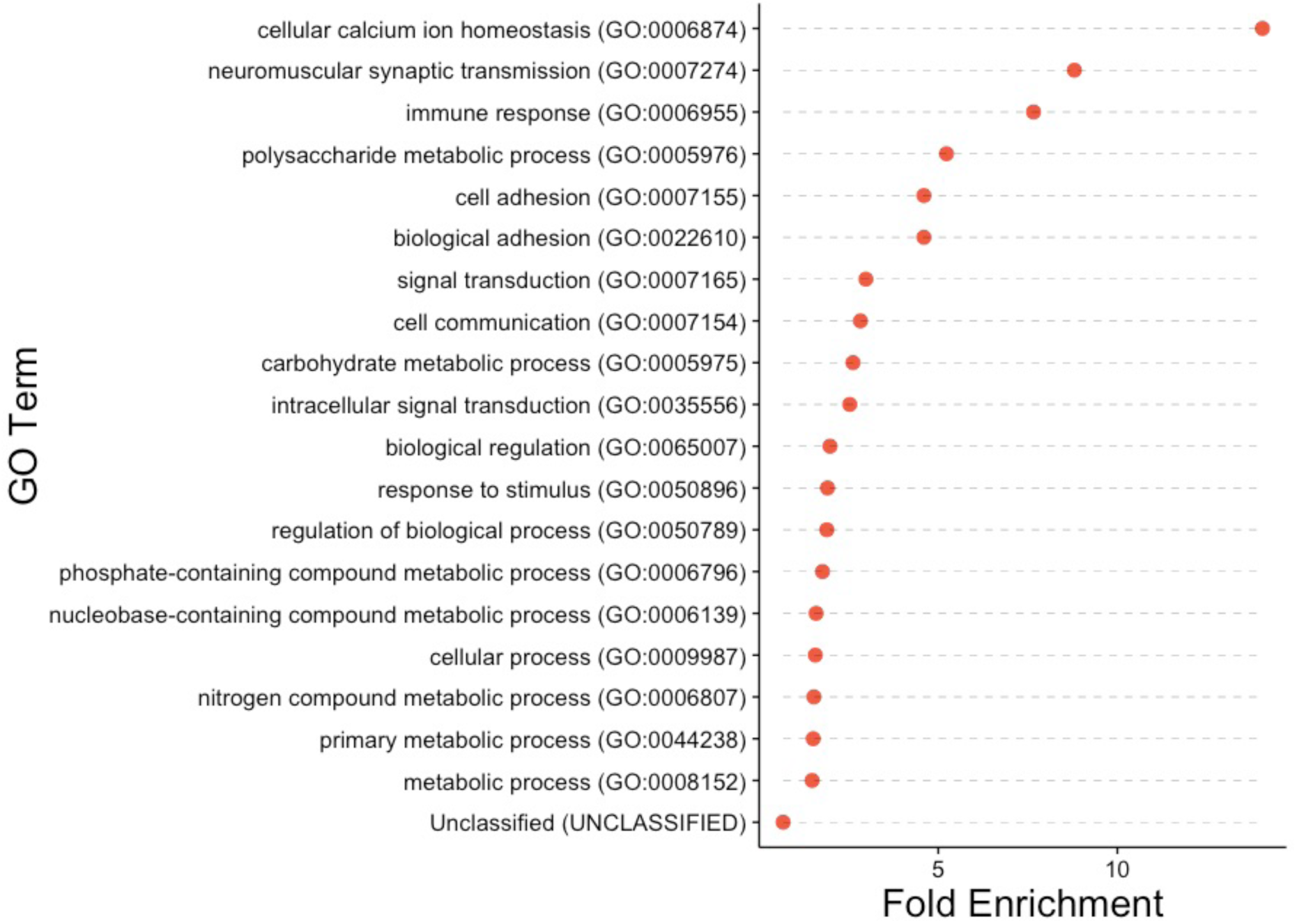
Gene ontology terms significantly overrepresented in the SNPs associated with PiGV resistance in larvae assayed on a common garden diet.

### Diet-specific resistance mechanisms

Our phenotypic data provided strong evidence that resistance evolves in an environmentally dependent manner, suggesting distinct genomic routes to resistance under specific nutritional conditions. This was particularly the case when we examined populations evolved on a nutrient limited diet but assayed on a high resource, common garden diet. In this situation, control populations exhibited much lower resistance than exposed populations. To identify the mechanism that provided this difference in resistance we repeated the association test for this subset of populations. Again, we found a large number of SNPs strongly correlated with resistance, suggesting a polygenic trait. Enrichment analyses were used to categorise these markers into biologically meaningful processes. In this case, different pathways were over-represented compared to the previous analysis that compared resistance on each population’s respective selection diet. For populations selected on a low nutrition diet but assayed on the common garden diet, cytoskeleton organization had a 5-fold enrichment followed by signal transduction and cell communication (Fig. S1). As a further demonstrative example, we investigated a scaffold that contained a high density of SNPs associated with PiGV resistance under these conditions (Fig. 5). After running a blast search on all genes on this scaffold we found a combination of developmental and metabolic genes interspersed with genes linked to viral and innate immunity as well as apoptosis (Table S2). Such linkage is suggestive of a pleiotropic effect or correlated selection, either of which could lead to the trade-off between immunity and development we observe.

**Figure 5.**
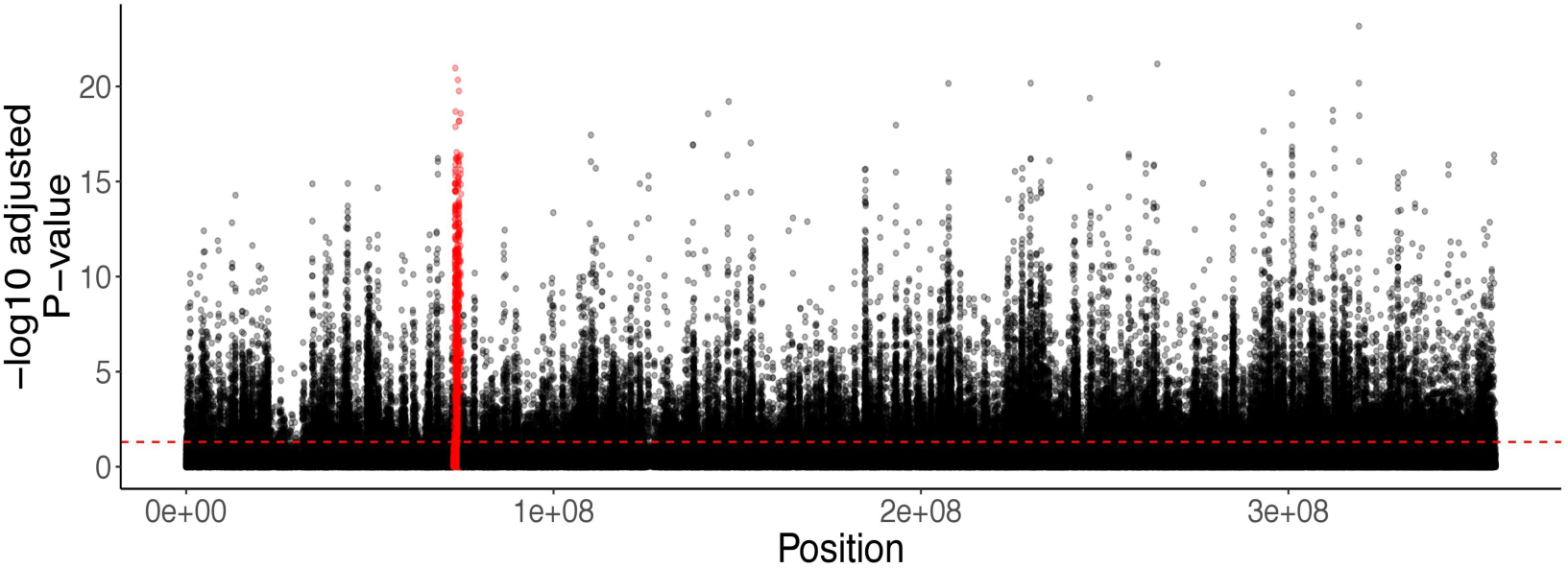
Manhattan plot of SNPs associated with resistance to PiGV on a low nutrition diet. Scaffold 23 is highlighted in red due to its high density of significantly associated SNPs. The putative functions of these genes are available in Table S2.

### PiGV resistance is a polygenic trait

The large number of SNPs associated with resistance in both the common garden and low nutrition diets suggests that resistance to PiGV infection is likely to be a highly polygenic trait. However, it is possible that the SNPs we associated with disease resistance are located in similar genomic regions, only appearing disparate due to the length of the scaffolds in our draft assembly. When we further assessed the proximity of SNPs, using improved scaffold lengths (increased from 0.5 to 5Mbp on average) from proximity-ligation sequencing we found many independent peaks of selection along the larger scaffolds (Fig. S2). This suggests that the genomic location of SNPs is not an artefact of the many small scaffolds that make up the assembly and that PiGV resistance is indeed a highly polygenic trait.

### Larval populations show little underlying population structure

Genome-wide association studies can lead to spurious correlations as a result of underlying population structure, where associations are a result of shared demographic history rather than a signature of selection. Whilst our populations were all derived from a single ancestral population, that had been out crossed repeatedly prior to the selection experiment, it is possible that the observed differences in allele frequency were the result of an underlying population structure. To rule this out we used a Bayesian approach to identity population structure naively on independent subsets of the SNP data. We identified very weak population structure suggesting that our results are unlikely to be spurious and found no clustering of populations that would be indicative underlying population structure (Fig. S3). We also found high reproducibility of results independent of which subset was used, suggesting our methods were robust (FMD always < 0.6, see Gautier 2015).

## Discussion

We used an evolve and re-sequence experiment (Schlötterer, Kofler, Versace, Tobler, & Franssen, 2015) to identify the resource dependence of the genetic basis of resistance to an insect DNA virus in an insect model system. We demonstrate that the evolution of resistance is diet specific, with populations selected on a nutrient limited diet utilising a different form of immunity. Surprisingly, we found no increase in resistance to PiGV exposure in populations selected on a high resource diet. In contrast, we found significantly higher survival in exposed populations selected on a low resource diet relative to controls. However, this effect was only apparent when such populations were assayed on a common garden diet, not when assayed on their low nutrition diet. Taken together, these results suggest that the resistance mechanism employed on low food is likely more complex than our bioassay can detect. For example, selection is acting on all life-stages during the selection experiment, but our assay by necessity only tests resistance at the ^3rd^ instar stage. This difference in resistance suggests that exposed populations selected on a low nutrient diet have evolved a low nutrition-specific resistance mechanism that is ineffective when reared and assayed on the common garden diet.

We also observed a trade-off between resistance and growth rate in nutrient limited populations. These results suggest that low nutrition environments may broadly limit the evolution of effective immune mechanisms. Whilst this nutrition-specific trade-off has been demonstrated before in two separate selection experiments (Boots & Begon, 1993; Boots, 2011) and shown to be genetic by comparison amongst inbred lines (Bartlett, Wilfert, & Boots, 2018), ours is the first study to use whole-genome resequencing to identify the genetic basis of such resistance. This demonstrates the genomic basis of one of the most well characterized genotypic trade-offs that shapes resistance to pathogens.

Our genome scan for resistance of all populations, regardless of selection diet, identified a number of biological processes involved in resistance to PiGV. A well-established phenomenon in baculovirus infections is apoptosis following cellular infection (reviewed in Schultz & Friesen 2009; Rohrmann 2013) and is thought to be a key immune response given that many baculoviruses encode anti-apoptosis genes (Clem, Hardwick, & Miller, 1996), including PiGV (Harrison, Rowley, & Funk, 2016). Although we did not identify apoptosis genes directly, the most overrepresented biological process was calcium ion homeostasis, which has previously been linked with baculovirus induced apoptosis (Xiu, Peng, & Hong, 2005). Other overrepresented biological processes include immune response and many metabolic and regulatory processes. This is understandable given that after the rapid global shutdown of mRNA and protein production following baculovirus infection, energy metabolism escapes this shutdown (Nguyen, Nielsen, & Reid, 2013). As such, our findings are broadly in line with existing mechanisms described in high nutrition diet.

Interestingly, when we picked out a number of candidate SNPs for further investigation (Fig. 3), we identified some potential targets for selection. Notably, SNP A (Fig. 3) encodes a mediator of RNA polymerase II transcription. This is of interest as all baculoviruses studied to date carry a protein that negatively regulates RNA polymerase II and is therefore likely to be a crucial component of baculovirus infectivity (Nguyen et al., 2013). We also identified a sodium and chloride dependent GABA transporter and a glycine receptor subunit, but it is difficult to link these to any specific baculovirus infection pathways. Finally, 3 of the candidates we selected were hypothetical proteins or in non-coding regions. This is potentially a result of working with a non-model organism for which genomic information is limited and may warrant further investigation.

As we identified a different phenotypic response in those populations selected on a low nutrition diet, we repeated the genome scan on just these populations to identify the putative resistance mechanism. In this instance, the most overrepresented biological process was cytoskeleton organization. This is interesting as baculoviruses are thought to manipulate the actin cytoskeleton during nucleocapsid transport and this is vital for successful infection and replication (Volkman, 2007). Baculoviruses use a cytoskeletal component (actin filaments) to both reach the nucleus and for transmission from the nucleus following nucleocapsid production (Marek, Merten, Galibert, Vlak, & van Oers, 2011; Ohkawa, Volkman, & Welch, 2010). As such, there is likely to be strong selection on resisting such manipulations. Other overrepresented biological processes included signal transduction, cell communication and cellular process. Furthermore, when we selected a single scaffold (scaffold 23) that was highly correlated with resistance in a low nutrition environment, we found it includes genes for innate immunity, intracellular virus transport, apoptosis and development (GO terms, see table S1). The close genetic linkage of such genes goes some way to explain the trade-off between resistance and development we observed. Finally, the disparity we see between resistance on a high nutrition diet vs. a nutrient limited diet suggests that under nutrient limitation, insects are forced to invest in intracellular resistance mechanisms that are traded off against growth rate. This is reinforced by our development data, where we find a much weaker trade-off when the same nutrient limited populations are assayed on a high-quality diet.

The number of SNPs correlated with observed resistance suggests that resistance is highly polygenic. To verify this, we used Hi-C scaffolding to improve scaffold lengths and still found many independent regions correlated with resistance. Taken together, this suggests that that resistance on both diets is a highly polygenic trait. A number of complex traits have previously been found to be highly polygenic in insects (Jha et al., 2015; Kang, Aggarwal, Rashkovetsky, Korol, & Michalak, 2016), suggesting that the mechanism of resistance in our system may also be complex, rather than a small number of typical immune genes as is typical in RNA virus immunity (Magwire et al., 2012). A recent study reports that a complex genetic architecture of many interacting genes can lead to genetic redundancy, with many competing beneficial alleles thus allowing rapid evolutionary responses. As such, it may be adaptive to rely on the complexity of polygenic traits (Barghi et al., 2018).

We have demonstrated that adaptation to a low nutrient diet can have profound effects on the underlying genetic architecture of virus resistance. We highlight a potential trade-off at the molecular level and describe putative resistance mechanisms that vary by diet. Our pool-seq approach has allowed a high level of replication at the population level and provided insights into the genetic nature of resistance. Further work will be required to fully characterize these mechanisms and functional validation of mutants in genome edited insects may soon be possible. Our results have implications for understanding wild insect populations and more broadly the role of nutrition across environments on pathogen resistance.

## Acknowledgements

SM and KR were supported by a NERC (UK) grant NE/J009784/1 awarded to MB and SP.

## Data Availability

The data that support the findings of this study will be openly available in DataDryad prior to publication. Sequence data is available from the European Bioinformatics Institute (EBI), under accession number PRJEB27964.

