## Supplemental Information for "The genomic basis of evolved virus resistance is dependent on environmental resources"

**Supplementary Information**

Table S1:

Top BLAST hits from all annotated genes on scaffold 23, a genomic region strongly associated with PiGV resistance in larvae selected and assayed on a low nutrition diet.

| Predicted Gene | *Plodia* Annotation ID | Uniprot GO Term | Organism |
| --- | --- | --- | --- |
| pre-mRNA-processing factor 17 isoform X1 | augustus_masked-scaffold23-abinit-gene-0.20 | mRNA processing, mRNA splicing | Amyelois transitella |
| nucleoporin NUP188 homolog | augustus_masked-scaffold23-abinit-gene-1.36 | Intracellular virus transport, viral transcription, constituent of nuclear pore | Amyelois transitella |
| tyrosine-protein kinase Drl | augustus_masked-scaffold23-abinit-gene-1.46 | Development protein, Kinase, Receptor | Amyelois transitella |
| RNA-binding protein 24-B-like isoform X1 | augustus_masked-scaffold23-abinit-gene-1.61 | RNA-binding | Helicoverpa armigera |
| lysine-specific demethylase 6A isoform X1 | augustus_masked-scaffold23-abinit-gene-2.23 | Cardiovascular system development, multicellular organism growth, respiratory system process | Amyelois transitella |
| toll-like receptor 8 | augustus_masked-scaffold23-abinit-gene-2.33 | Innate immunity, inflammatory response | Amyelois transitella |
| sodium-coupled monocarboxylate transporter 1-like | augustus_masked-scaffold23-abinit-gene-3.30 | Apoptosis, Ion transport, Sodium transport | Amyelois transitella |
| carbonic anhydrase 7 | maker-scaffold23-augustus-gene-0.43 | Lyase, one-carbon metabolic process, regulation of synaptic transmission | Amyelois transitella |
| uncharacterized phosphotransferase YvkC-like | maker-scaffold23-augustus-gene-1.102 | ATP binding, kinase activity | Amyelois transitella |
| cleft lip and palate transmembrane protein 1-like protein | maker-scaffold23-augustus-gene-1.104 | Cell differentiation, multicellular organism development, | Amyelois transitella |
| PAS domain-containing serine/threonine-protein kinase | maker-scaffold23-augustus-gene-1.74 | ATP binding, regulation of energy homeostasis | Amyelois transitella |
| neurofilament medium polypeptide-like isoform X2 | maker-scaffold23-augustus-gene-1.77 | Microtubule binding, cytoskeleton constituent, axon development | Amyelois transitella |
| double-stranded RNA-binding protein Staufen homolog 2 | maker-scaffold23-augustus-gene-1.90 | Double stranded RNA-binding, Transport | Spodoptera litura |
| hypothetical protein KGM_201520 | maker-scaffold23-augustus-gene-1.92 | NA | Danaus plexippus plexippus |
| guanine nucleotide exchange factor DBS-like isoform X3 | maker-scaffold23-augustus-gene-1.94 | Signaling pathways, positive regulator of apoptosis | Amyelois transitella |
| mediator of RNA polymerase II transcription subunit 30 isoform X1 | maker-scaffold23-augustus-gene-1.99 | Signaling pathways | Bombyx mori |
| PREDICTED: uncharacterized protein LOC106138209 | maker-scaffold23-augustus-gene-2.50 | NA | Amyelois transitella |
| PREDICTED: sequestosome-1-like isoform X2 | maker-scaffold23-augustus-gene-2.54 | Apoptosis, Autophagy, Differentiation, Immunity | Amyelois transitella |
| PREDICTED: sodium-coupled monocarboxylate transporter 1-like | maker-scaffold23-augustus-gene-2.57 | Apoptosis, Ion transport, Sodium Transport | Amyelois transitella |
| PREDICTED: dual specificity protein phosphatase 13 isoform B-like isoform X2 | maker-scaffold23-augustus-gene-2.59 | Hydrolase, protein phosphatase | Amyelois transitella |
| PREDICTED: SHC-transforming protein 1 isoform X1 | maker-scaffold23-augustus-gene-2.60 | Angiogenesis, Growth regulation, Host-virus interaction | Amyelois transitella |
| PREDICTED: arf-GAP with SH3 domain, ANK repeat and PH domain-containing protein 2 | maker-scaffold23-augustus-gene-2.62 | GTPase activation | Amyelois transitella |
| PREDICTED: exosome complex component RRP42 | maker-scaffold23-augustus-gene-2.63 | rRNA-binding, rRNA processing | Amyelois transitella] |
| PREDICTED: pancreatic triacylglycerol lipase-like | maker-scaffold23-augustus-gene-3.58 | Lipid metabolism | Amyelois transitella |
| N-alpha-acetyltransferase 35, NatC auxiliary subunit isoform X1 | maker-scaffold23-augustus-gene-3.67 | Apoptosis, Muscle cell proliferation | Helicoverpa armigera |
| PREDICTED: pancreatic triacylglycerol lipase-like | maker-scaffold23-augustus-gene-3.71 | Lipid metabolism | Amyelois transitella |
| PREDICTED: adenylate cyclase type 7-like isoform X3 | maker-scaffold23-augustus-gene-3.75 | cAMP biosynthesis, signal transduction | Amyelois transitella |
| PREDICTED: molybdenum cofactor sulfurase | maker-scaffold23-augustus-gene-3.77 | Transferase | Amyelois transitella |
| PREDICTED: poly(rC)-binding protein 2 isoform X1 | maker-scaffold23-augustus-gene-4.66 | Antiviral defence, Immunity, Innate Immunity, Viral RNA replication | Neodiprion lecontei |
| PREDICTED: arylsulfatase B-like | maker-scaffold23-augustus-gene-4.72 | Development pathways and metabolic processes | Amyelois transitella |
| PREDICTED: brachyurin-like | maker-scaffold23-augustus-gene-4.79 | Collagen degradation | Amyelois transitella |

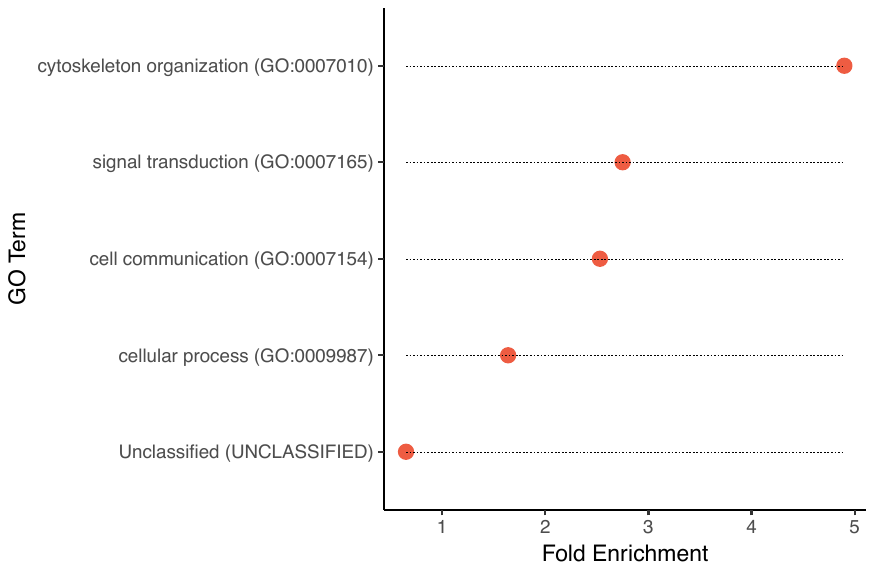

Figure S1.

Overrepresented gene ontology terms based on all SNPs significantly associated with PiGV resistance on a common garden diet for populations selected on a low nutrition diet.

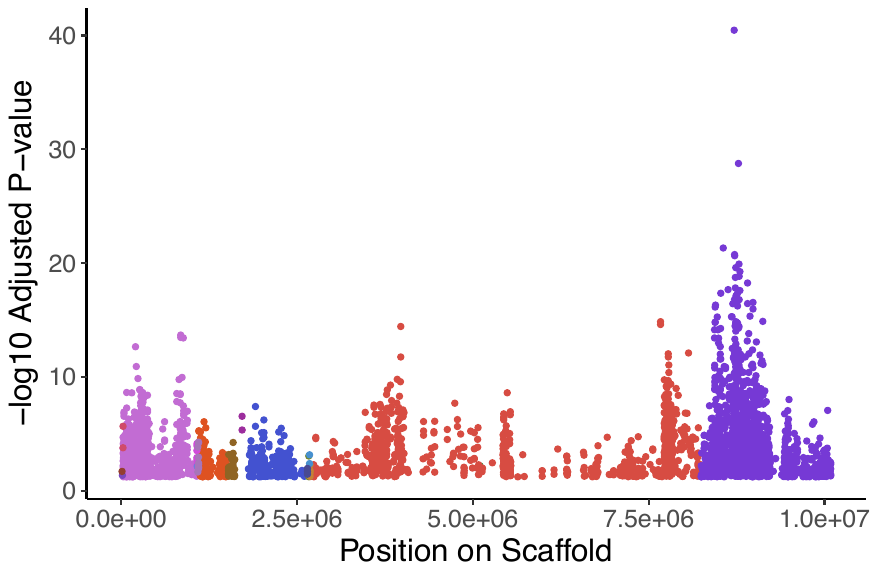

Figure S2.

Improved scaffold resolution based on proximity sequencing. Each color represents an independent scaffold from the original genome assembly, showing their relative locations based on the resolved scaffold.

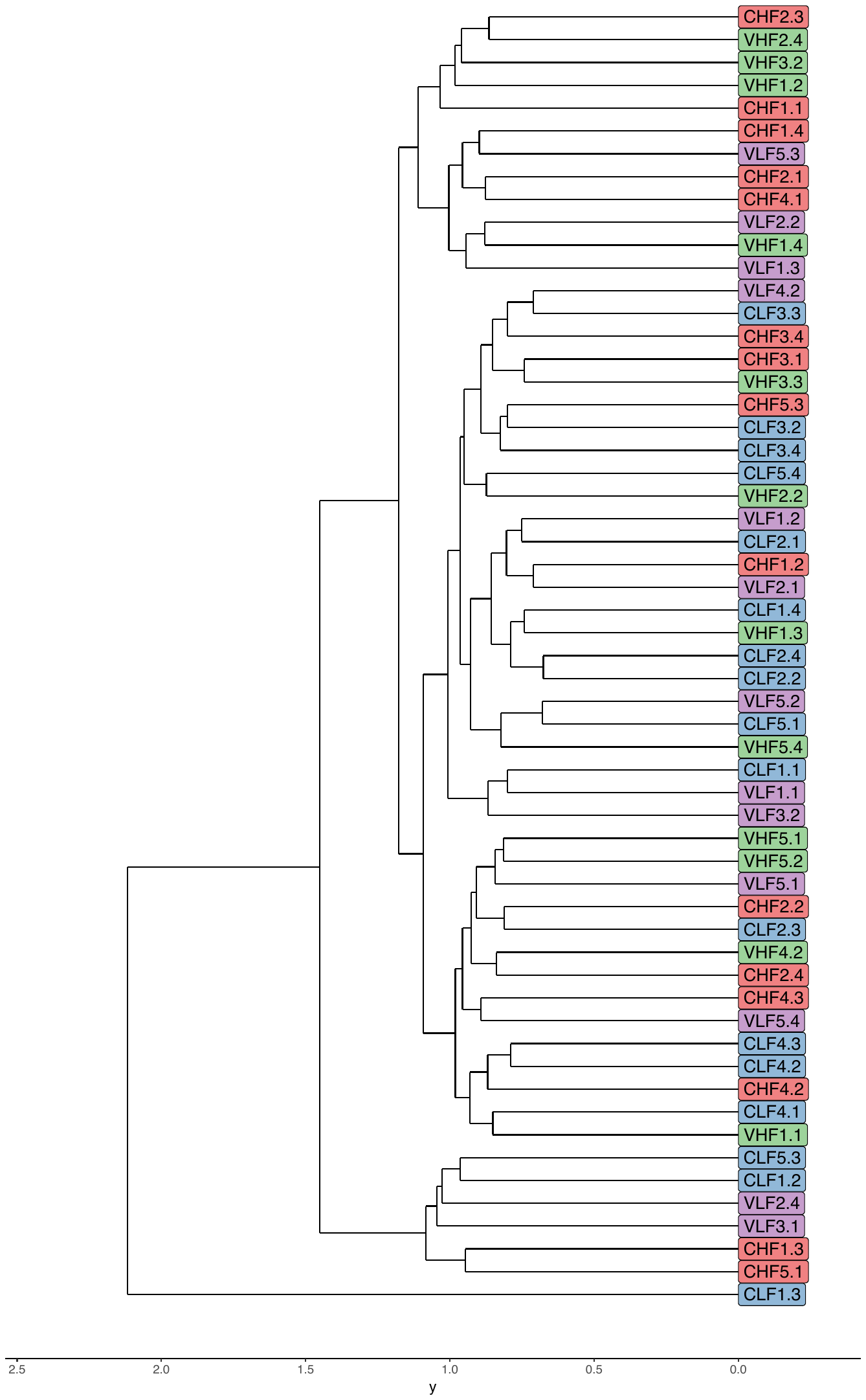

Figure S3

Hierarchical clustering of populations based on a naïve Bayesian genome-wide scan for underlying population structure. Colors denote selection treatment groups. C = Control, V = Virus, H = High nutrition, L = Low nutrition.
